# Mitochondrial morphology and function are tuned by microtubule association

**DOI:** 10.64898/2026.06.13.731791

**Authors:** Nida Ul Fatima, Gregory MI Redpath, Zeenat Jahan, Olivia Nassar, Nicholas Ariotti, Vaishnavi Ananthanarayanan

**Affiliations:** EMBL Australia Node in Single Molecule Science, Department of Molecular Medicine, School of Biomedical Sciences, University of New South Wales, Sydney, NSW 2052, Australia; Motor Neuron Disease Research Centre, Macquarie Medical School, Macquarie University, Sydney, NSW 2113, Australia; Chris O’Brien Lifehouse, Sydney, NSW 2050, Australia; Institute for Molecular Bioscience, The University of Queensland, St Lucia, Queensland 4072, Australia

## Abstract

Mitochondrial morphology is regulated through dynamic fission and fusion processes that are critical for cellular metabolism and homeostasis. Microtubules and mitochondria are functionally coupled across many cellular contexts, yet the mechanistic basis of this relationship remains poorly understood. Here, we reveal that microtubule association is a key regulator of mitochondrial morphology and function. Pharmacological or genetic manipulation of microtubule dynamics controls mitochondrial membrane potential and respiratory capacity, with stabilisation driving elongation, and depolymerisation triggering Drp1-dependent fission and reduced respiration. Strikingly, we identify increased microtubule polymerisation as a hallmark of starvation-induced mitochondrial hyperfusion, a key adaptive response to nutrient stress, and show that this expanded microtubule network is an essential prerequisite for the starvation response. We demonstrate that this regulation operates independently of canonical tubulin post-translational modifications that stabilise microtubules. Rather, increased mitochondria-microtubule association, driven by network expansion, is both necessary and sufficient to drive mitochondrial hyperfusion, as demonstrated by forced mitochondria-microtubule tethering. Taken together, our findings establish microtubule binding as a key upstream regulator of mitochondrial network dynamics, with direct implication for understanding how cells coordinate cytoskeletal organisation with metabolic adaptation during physiological stress.

## Introduction

Mitochondria are dynamic organelles that continuously undergo fission and fusion, with the equilibrium between fission and fusion determining mitochondrial morphology^1,2^. Mitochondrial fission and fusion are driven by GTPases. Drp1 is the key mitochondrial fission protein^3^, and is recruited to the mitochondrial outer membrane by MiD49, MiD51 and Mff^4,5^. Mfn1/Mfn2 and Opa1 mediate outer and inner mitochondrial membrane fusion respectively^6,7^. While the central players that control mitochondrial fission and fusion have been identified, the cellular cues that dictate the spatiotemporal localisation and assembly of these fission and fusion proteins in physiological conditions remain understudied.

We recently showed a strong link between mitochondrial morphology and microtubule (MT) length in fission yeast cells (*Schizosaccharomyces pombe*)^8–10^. Like in mammalian cells, mitochondria associate with microtubules in *S. pombe*^11^. However, unlike in mammalian cells, *S. pombe* mitochondria are not transported on MTs by motor proteins, but rather only use MTs as physical scaffolds^12^. In *S. pombe*, MT binding reduces mitochondrial fission frequency due to inability of Dnm1 (yeast Drp1) to assemble around mitochondria when they were bound to MTs^8,9^. As a result, cells with short MTs (with less mitochondria-binding scaffolds) had more mitochondrial fission and more fragmented mitochondria, whereas cells with longer MTs had longer mitochondria due to reduced mitochondrial fission^8,10^. It is unknown how mitochondrial morphology might change with perturbation of MT lengths and stability in mammalian cells. However, it is tempting to speculate a strong association between MTs and mitochondrial dynamics in mammalian cells given that about 70% of mitochondria remain stationary on MTs even in cells with high energy demands such as neurons^13^.

MTs serve to anchor mitochondria along the axons of neurons to enable uniform energy distribution along these long cells. For instance, syntaphilin^13,14^ has the ability to dock mitochondria along neuronal MTs in a Ca^2+^-depedent manner, but is expressed primarily in neurons and immune cells. Mitochondria link to MTs via motor proteins cytoplasmic dynein and kinesin for transport in neurons and other cell types^15^. Motor adaptor proteins TRAK1 and TRAK2 associate with Miro1/2 on the mitochondrial outer membrane to bridge MTs to mitochondria^16,17^. While motors and their adaptors participate in moving mitochondria on MTs, a ubiquitous protein linker that mediates persistent docking of mitochondria on MTs across all cell types has not yet been identified.

Appropriate mitochondrial morphology is not only necessary to enable key processes such as mitochondrial inheritance, partitioning and mitophagy, but is also linked to mitochondrial function. Elongated mitochondrial networks are associated with increased oxidative phosphorylation and mitochondrial membrane potential, metabolic adaptation, and reduced reactive oxygen species (ROS) production, whereas fragmented mitochondria typically display lowered mitochondrial membrane potential with simultaneous increase in ROS^18^.

In this work, we employed a combination of pharmacological and physiological perturbations, fission and fusion protein knockouts, and high resolution light and electron microscopy to investigate the role of MT association in dictating mitochondrial morphology, how this relates to mitochondrial function in normal and starved cells, and show that forced association of mitochondria to MTs is sufficient to induce mitochondrial hyperfusion. Taken together, this work establishes microtubule association as an upstream regulator of mitochondrial morphology and function.

## Materials and methods

### Cell culture

Wild-type HeLa cells (ATCC CCL-2, RRID: CVCL_0030) and HEK293 cells (ATCC: CRL-1573) were maintained in Dulbecco’s Modified Eagle Medium (DMEM) supplemented with 10% fetal bovine serum, 100 U/mL penicillin, 0.1 mg/mL streptomycin, and 2 mM L-glutamine (Cat# AT066, HiMedia Laboratories; Cat# 11965092, Thermo Fisher Scientific). Cells were cultured at 37°C in a humidified 5% CO₂ incubator.

### Cell transfection

Cells were plated in glass-bottom dishes (Ibidi, Cat# 81518) or 18 well chamber slides (Ibidi, Cat# 81816) and transfected overnight with plasmids including ɑ-tubulin-GFP (Addgene plasmid 56450, a gift from Michael Davidson), PA-GFP (generous gift from Prof Deepak Saini, Indian Institute of Science, Bengaluru), Tom20-mCherry-FKBP (Addgene plasmid 162436, a gift from Takanari Inoue) and EMTB-ECFP-FRB (a generous gift from Prof. Yu-Chun Lin, National Ting Hua University) using Lipofectamine 3000 (Thermo Fisher Scientific, L3000001) according to the manufacturer’s instructions.

### Depletion of MAPs with RNAi

MAP1S Pre-designed Silencer Select siRNA was purchased from Thermo Fisher Scientific (# s30424). MAP4S siRNA Duplexes were purchased from Sigma Aldrich (#VC30002). siRNA was transfected into cells using a modified Lipofectamine 3000 protocol with 75 pmol siRNA and 6 uL Lipofectamine 3000 reagent in 250 uL OptiMEM per well of a 6 well plate. Cells were grown for 48 hours following siRNA transfection before analysis.

### Drp1^ko^ and Opa1^ko^ construction

Knockout of Drp1 and Opa1 was performed using CRISPR/Cas9. Cas9 guides targeting Drp1 (GAAGATAAACGGAAAACAA, TTCAGCTGTATCACGAGACA) and Opa1 (CGCTTTATGACAGAACCGAA, TTACCCGGTCAAGTTCCCGC) were purchased from IDT and cloned into spCas9-2A-GFP (Addgene plasmid number 48138, a gift from Feng Zhang) using BbsI-HF (NEB #R3539). HeLa cells were transfected with the guide-containing Cas9 plasmids using Lipofectamine 3000 and single cell sorting for GFP-positive cells undertaken by staff at the UNSW Flow Cytometry Facility. Single cell clones were expanded prior to analysis for protein expression by Western blot.

### Drug treatments

For live-cell imaging, HeLa and HEK293 cells were seeded in glass-bottom dishes and treated with either 10 µM nocodazole (Sigma Aldrich M1404) or 1 µM paclitaxel (Sigma Aldrich T7191) for 30 minutes. DMSO-treated cells served as controls. For rapamycin experiments, cells were exposed to 200 nM rapamycin (Abcam ab120224) for 1 h prior to imaging.

### Inhibitor and dye treatments

Mitochondria were labelled using 100 nM MitoTracker Deep Red (Thermo Fisher Scientific, M22426) following the manufacturer’s protocol. For drug-treatment conditions, staining was performed during the final 20 min of drug exposure. For imaging the mitochondrial membrane potential, cells were incubated with 100 nM TMRE (Thermo Fisher Scientific, T669) for 30 min, washed, and imaged using the 560-nm channel. To inhibit ROCK, cells were treated with 10 µM Y-27632 (MedChem Express, HY-10071) for 30 min, washed, and imaged.

### Immunostaining

Cells were fixed either with 4% paraformaldehyde (20 min at room temperature) followed by permeabilisation with 0.4% Triton X-100 in PBS (15 min), or with ice-cold methanol (15 min on ice). After three PBS washes, cells were blocked in 5% BSA for 30 min, incubated with primary antibodies overnight, washed, and incubated with secondary antibodies for 1 h prior to imaging.

### Primary antibodies used

- MAP1S (Abcam, ab254930; rabbit polyclonal), 1:50
- MAP4 (Abcam, ab224543; rabbit polyclonal), 1:1000
- Opa1 (CST, 67589; rabbit monoclonal), 1:1000
- Drp1 (Proteintech, 12957-1-AP; Rabbit polyclonal), 1:5000
- Tom20 (Thermo Fisher, MA5-34964; rabbit monoclonal), 1:100
- Detyrosinylated α-Tubulin (Abcam, ab48389; rabbit polyclonal), 1:200
- Tyrosinylated α-Tubulin (Abcam, ab6160; rat monoclonal), 1:5000
- Acetylated tubulin (Sigma Aldrich, T7451; mouse monoclonal), 1:1000
- â-Tubulin-405 (Abcam, EPR16774; rabbit monoclonal), 1:2000

### Microscopy

Confocal imaging was performed using a Nikon A1 or Nikon AX-R2 microscope equipped with a 60× 1.20 NA water-immersion objective, or a Nikon Ti2 with Yokogawa W1 spinning disc confocal with a 100x 1.4 NA oil-immersion objective. Z-stacks were acquired at 1-µm intervals. For PA-GFP experiments, circular ROI of 1.68 µm diameter was drawn and stimulated with a 405nm laser for 1s at 5% laser power and imaged 3s after stimulation. Live-cell imaging was performed at 37°C in imaging buffer (140 mM NaCl, 2.5 mM KCl, 1.8 mM CaCl₂, 1 mM MgCl₂, 4 mg/mL D-glucose, and 20 mM HEPES). Fixed-cell imaging was conducted at room temperature (23°C) in PBS. Imaging was carried out at the Katharina Gaus Light Microscopy Facility, Mark Wainwright Analytical Centre, UNSW.

### Transmission electron microscopy

HeLa cells were fixed for electron microscopy in 2.5% glutaraldehyde (Sigma-Aldrich, G5882) in PBS (pH 7.4) for 1 hour at 37°C. Fixed cells were washed 3 times in PBS, then post-fixed in 2% OsO4 with 1.5% potassium ferricyanide. Samples were then washed in double-distilled H2O (ddH_2_O), incubated in a 1% (w/v) thiocarbohydrazide solution and postfixed again in 2% OsO4. Cells were washed (ddH_2_O) and stained en bloc with 1% uranyl acetate and subsequently with lead aspartate solution (20 mM lead nitrate, 30 mM aspartic acid, pH 5.5). Cells were then serially infiltrated with LX112 resin and polymerised for 48 h. Ultrathin sections were cut on an ultramicrotome (UC6: Leica) and imaged at 80 kV on a Hitachi 7700 for low magnification analyses of mitochondrial shape and a JEOL JEM-1400Plus TEM fitted with a Phurona EMSIS CMOS camera for high-magnification analyses. Segmentation for mitochondrial size and shape analyses was performed with the Segment Anything SAMJ plugin in Fiji. Imaging and analyses were performed in a blinded fashion.

### Serial block face scanning electron microscopy (SBF-SEM)

Samples were fixed, stained, and resin infiltrated as above. Following, SBF-SEM imaging was performed in an ApreoVS scanning electron microscope (Thermofisher Scientific). Polymerised LX112 resin was thinned and different experimental conditions were sandwiched together for vertically oriented slicing with additional LX112 resin (i.e., DMSO-control and paclitaxel-treated cells in the same block). Low magnification tiled (2x2) overviews were acquired at every 10 slices to map emerging cells for higher-resolution imaging. High-resolution imaging was performed using MAPS software (Thermofisher Scientific) at a pixel size of 10nm x10nm with dwell time of 2µs and a z-depth of 100nm. Tiled image stacks were assembled and aligned in Fiji and iteratively realigned in etomo (IMOD) (https://bio3d.colorado.edu/imod/). Final aligned data was denoised using BM3D^19^ (Oak Ridge National Laboratory, “bm3dornl: High-performance BM3D denoising for imaging,” GitHub repository, 2024, https://github.com/ornlneutronimaging/bm3dornl) and semi-automated segmentation was performed in IMOD with visualisation generated using ChimeraX10.1 (UCSF).

### Seahorse Assay

Oxygen Consumption Rate (OCR) was measured in intact, adherent cells using the Agilent Seahorse XF Cell Mito Stress Test Kit on an XFe96 Extracellular Flux Analyzer. HeLa cells were seeded one day prior to analysis at 3 × 10⁴ cells/well, in XF96 plates. Prior to the assay, cells were assessed for viability and confluency. Cells were washed in XF assay medium supplemented with 25 mM glucose, 1 mM sodium pyruvate, and 2 mM L-glutamine and treated with paclitaxel, nocodazole, or vehicle control before analysis. Basal mitochondrial respiration was recorded, followed by sequential injections of oligomycin A (1.5 μM), BAM15 (10 μM; concentration optimized in preliminary experiments), and rotenone/antimycin A (0.5 μM each). Four unseeded wells per plate served as background controls. Basal respiration, ATP-linked respiration, maximal respiration, and proton leak were calculated using standard Seahorse definitions. All conditions were measured in technical triplicate, and experiments were repeated independently three times.

### Starvation Assays

The starvation protocol was performed as described previously^20^. Briefly, 24h following plating, HeLa cells were either left in their original media (DMEM, unstarved), or washed twice with HBSS, then incubated in HBSS for 5 hours. Following the 5 hour starvation, cells were used for analysis. For trichostatin A treatments (TSA, Sigma-Aldrich T8552), starved and unstarved cells were treated with 200 nM at the start of the starvation time course.

### Image Analysis

Mitochondrial segmentation on 2D maximum intensity projections of fluorescence images was performed using the Mitochondria Analyzer pipeline^21^ in Fiji/ImageJ. The Mitochondria Analyzer plugin was employed in this work since it resulted in the most accurate mitochondrial morphologies in simulated and real mitochondrial images when compared to five other popular plugins in Fiji/ImageJ^22^. The mean area and the mean length of the major axis of all the mitochondria across a cell was used to report on the mitochondrial area and mitochondrial length plots. The mean for each repeat (with cell numbers listed in the figure captions for each data set) was calculated, and the mitochondrial area and length values were normalised to the mean value of the control. These repeat means were then subjected to statistical analysis as described below. For estimation of the intensity of tubulin or acetylated, tyrosinated, or detyrosinated tubulin, the integrated density and/or mean intensity per cell was calculated and normalised to the mean of the control, and the mean of the repeats were subjected to statistical analysis.

For manual classification of mitochondrial morphology (Fig. S1), individual cells were cropped from fluorescence images by one observer and saved into separate folders, which were subsequently blinded by a second observer to prevent classification bias. Manual classification of each cell image was then performed by the first observer on the blinded dataset. Cells were assigned to one of three morphological categories based on the appearance of the mitochondrial network: Category 1, comprising cells with predominantly elongated mitochondria; Category 2, comprising cells with a mixed population of elongated and fragmented mitochondria; and Category 3, comprising cells with predominantly fragmented mitochondria.

For presentation in the figures, individual cells were cropped, and the Fiji ‘Clear outside’ function was applied outside the region containing the cell of interest.

### Statistical analysis

All plots were generated in MATLAB (Mathworks). To test the statistical significance of the difference between distributions we used ordinary one-way ANOVA with Tukey Kramer post hoc test/Dunnett’s multiple comparisons test, or Kruskal Wallis test with Bonferroni correction for non-parametric data. The specific statistical tests used for each dataset are mentioned in the corresponding figure captions. The figures were organised and prepared in Adobe Illustrator.

## Results

### Nocodazole-induced microtubule depolymerisation results in mitochondrial fission, and paclitaxel-induced microtubule stabilisation leads to elongated mitochondrial networks

Previously, we reported that fission yeast mitochondria that were bound to microtubules were less likely to undergo fission due to the inhibition of the Dnm1 (yeast Drp1) assembly around mitochondria^8^. Alteration of MT dynamics by using chemical inhibitors, or by using genetic manipulation resulted in changed mitochondrial morphology, wherein cells with longer MT had longer mitochondria and vice versa. We first tested if a similar relationship between MT and mitochondrial form could be observed in mammalian cells. Upon treatment with the MT stabilising drug paclitaxel, mitochondria appeared more networked and elongated compared to DMSO-treated control HeLa cells, whereas cells treated with the MT-depolymerising drug nocodazole showed more fragmented mitochondria than control or paclitaxel-treated cells (Fig. 1A). Quantification of mitochondrial area and length mirrored these findings, with paclitaxel-treated cells displaying larger and longer mitochondria, whereas nocodazole-treated cells showing smaller and shorter mitochondria (Fig. 1B, C; Fig. S1A, B). Visualisation of mitochondria in paclitaxel- and nocodazole-treated cells in 2D EM thin sections reinforced this observation, with mitochondria in paclitaxel-treated cells exhibiting longer mitochondria with low circularity, and those in nocodazole-treated cells showing shorter, and slightly more circular mitochondria (Fig. 1 D, E). So too, volume EM (SBF-SEM) of paclitaxel-treated cells revealed increased mitochondrial volumes compared to DMSO-treated cells (Fig. S1C, D), confirming our light microscopy observations. We asked if this phenomenon was specific to HeLa cells or if it was reproducible in other mammalian cells in culture. We observed that HEK293 cells also displayed mitochondrial behavior consistent with our findings in HeLa cells upon treatment with paclitaxel or nocodazole (Fig. S1E-H).

**Figure 1.**
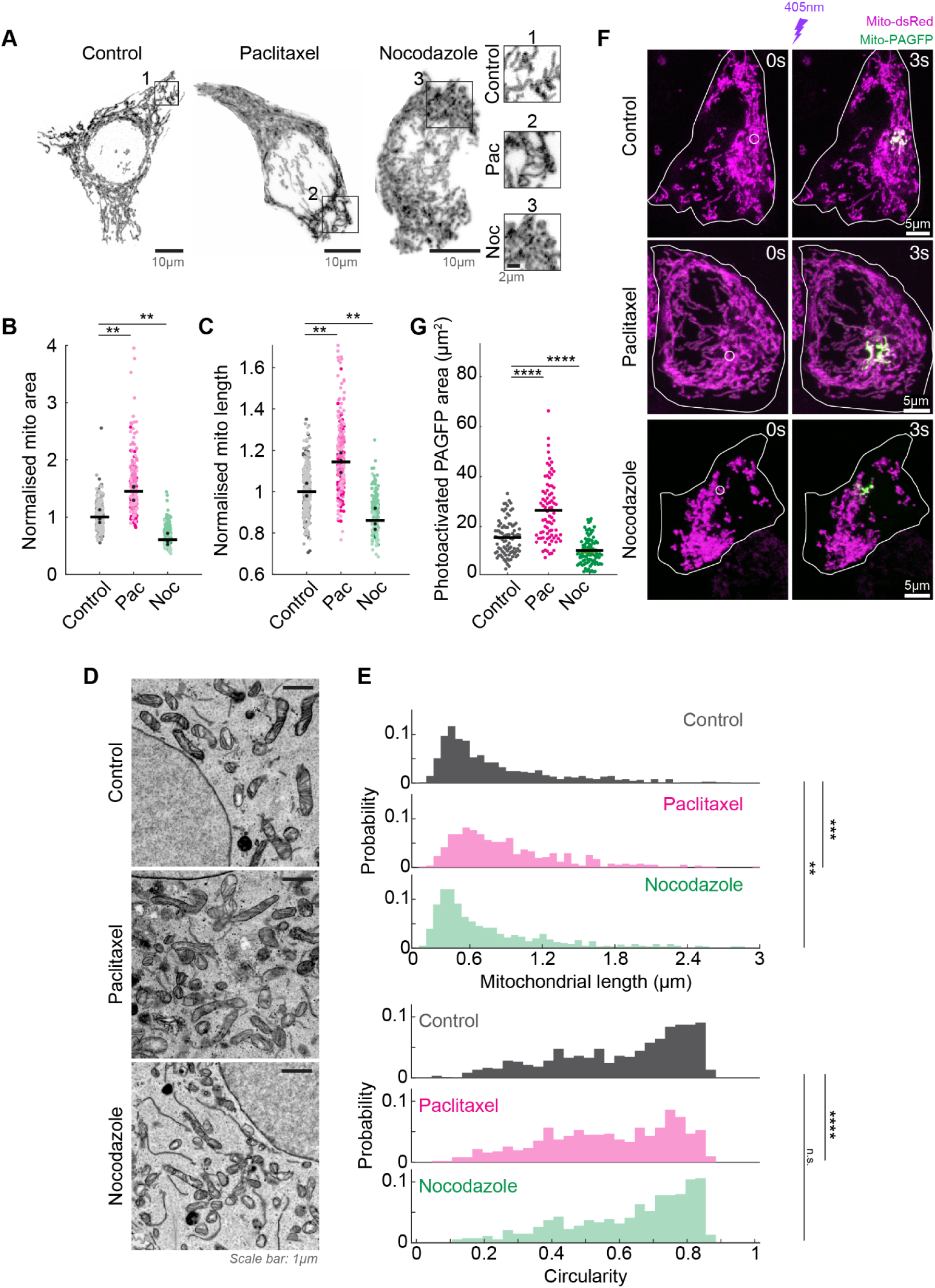
Microtubule perturbation alters mitochondrial form. **(A)** Representative live-cell confocal microscopy images of Mitotracker Deep Red-labelled mitochondria in HeLa cells following treatment with DMSO (control), paclitaxel (pac), and nocodazole (noc) for 1h. The enlarged versions of the regions marked 1, 2 and 3 from control, paclitaxel- and nocodazole-treated cells appear on the right. Scatter plots of **(B)** mitochondrial area and **(C)** mitochondrial lengths in control, paclitaxel- and nocodazole-treated cells normalised to the mean of the control cells. **(D)** TEM images of thin sections of cells treated with DMSO (control, top), paclitaxel (middle), and nocodazole (bottom) exhibiting different mitochondrial morphologies. **(E)** Histograms of mitochondrial length (top) and circularity (bottom) in control, paclitaxel and nocodazole-treated cells. Data were obtained from 2 repeats with >23 cells in each repeat. Asterisks represent significance: **p=0.002, ***p=0.0003, ****p=9.2x10^-7^, and ‘n.s.’ implies no significant difference, Kruskal Wallis test with Bonferroni’s correction. **(F)** Representative live-cell confocal microscopy images of control, paclitaxel- and nocodazole-treated cells transiently transfected with mito-dsRed (magenta) and mito-PAGFP (green) before (left, 0s) and after (right, 3s) photo stimulation with 405 nm in the region marked by the white circle. **(G)** Quantification of the photoactivated area with PAGFP fluorescence 3s after stimulation in control, paclitaxel- and nocodazole-treated cells. In **B**, **C** and **G**, data were obtained from N=3 independent repeats, with 30-100 cells in each repeat. The black circles represent the means of the individual repeats, and the black lines represent the mean of the 3 repeats. Asterisks (**) represent p<0.01, (****) represent p<0.0001, 1-way ANOVA with Tukey-Kramer post hoc test.

To confirm our results with an orthogonal technique, we assayed the mitochondrial network status in cells expressing mitochondria localized photoactivable (PA)-GFP. Mitochondrial matrix-localised PA-GFP has been used previously to report on mitochondrial fusion dynamics^23^: upon stimulation of PA-GFP in mitochondria with 405 nm laser in a small region of the cell, the fluorescence from mitochondrial (mt)-PA-GFP was visible within seconds in mitochondria fused to the original photoactivated region due to rapid diffusion of mt-PA-GFP through the connected matrices. The final photoactivation area was thus a readout of the network status of mitochondria, with a higher photoactivation area indicating more fused mitochondrial networks and a lower photoactivation area indicating a fragmented network. In paclitaxel-treated cells expressing mt-PA-GFP, we indeed observed a significantly higher photoactivation area compared to control cells, and nocodazole-treated cells showed photoactivation areas lower than both control and paclitaxel-treated cells (Fig. 1F, G). This confirmed our finding that mitochondria were more fused in cells with more stable, paclitaxel-treated MT networks, whereas cells without MTs had fragmented mitochondria.

### Mitochondrial fission upon microtubule depolymerisation is Drp1-dependent

Next, we asked if the reduced mitochondrial size in nocodazole-treated cells was the outcome of enhanced fission. We knocked out the primary mitochondrial fission protein Drp1 (Fig. S2A) and visualised the mitochondria in these cells in the presence and absence of nocodazole (Fig. 2A). Mitochondria in Drp1^ko^ cells treated with DMSO appeared elongated as expected due to uncontrolled fusion in the absence of mitochondrial fission (Fig. 2A-C). In the presence of nocodazole, while wild-type (WT) cells displayed fragmented mitochondria, Drp1^ko^ cells continued to show elongated mitochondrial networks (Fig. 2A-C), indicating that the fragmentation of mitochondria in the absence of MTs in these cells required Drp1-dependent fission.

**Figure 2.**
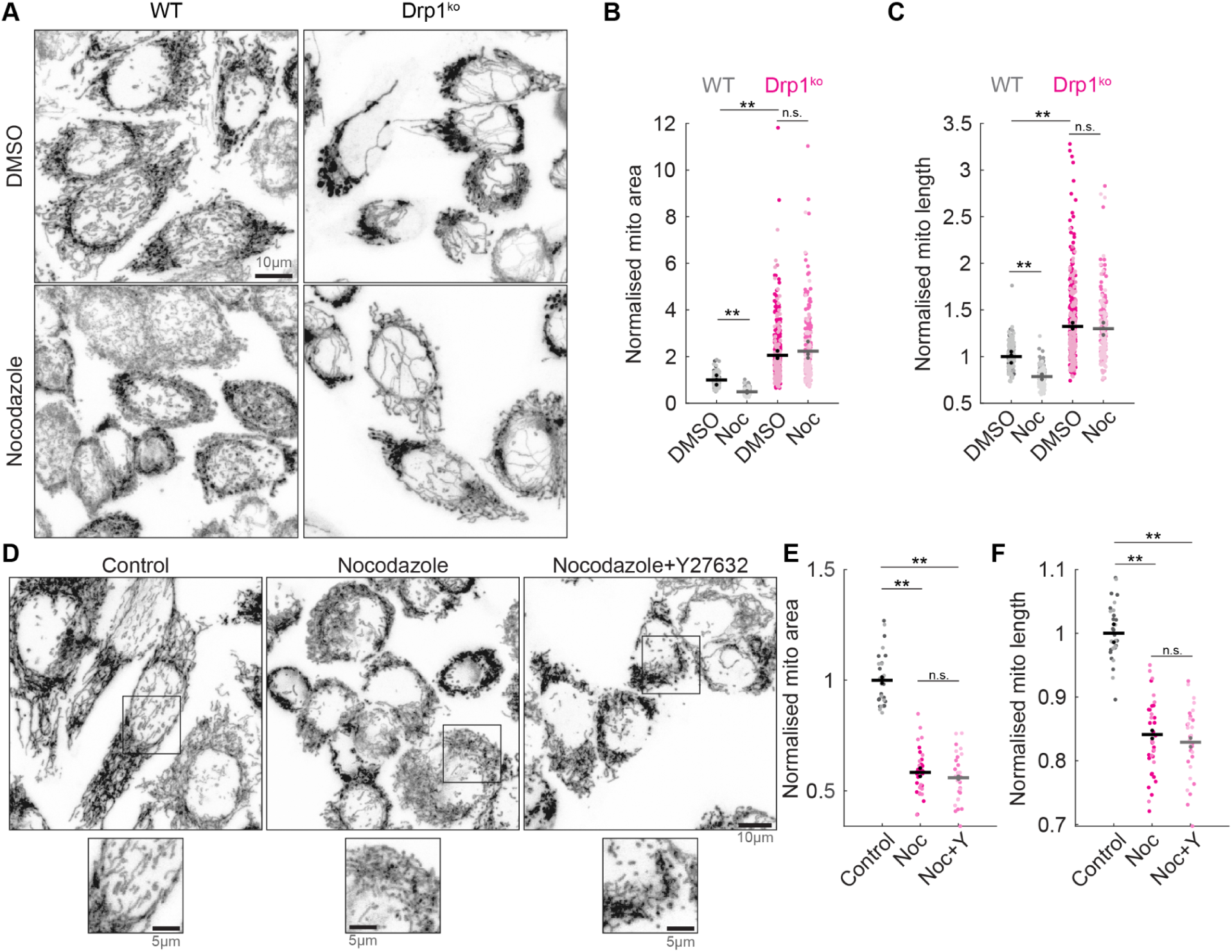
Increased microtubule fission upon nocodazole treatment is Drp1-dependent. **(A)** Representative live-cell confocal microscopy images of Mitotracker Deep Red-labelled mitochondria in wildtype (WT) or Drp1^ko^ HeLa cells following treatment with DMSO (top row), and nocodazole (bottom row) for 1h. Scatter plots of **(B)** mitochondrial area and **(C)** mitochondrial lengths in DMSO, and nocodazole-treated WT and Drp1^ko^ cells normalized to the mean of the WT DMSO-treated cells. **(D)** Representative live-cell confocal microscopy images of Mitotracker Deep Red-labelled mitochondria in HeLa cells following treatment with DMSO (control), nocodazole (noc), and nocodazole+Y-27632 (noc+Y) for 1h. The enlarged versions of the regions marked with the black squares from control, nocodazole-, and noc+Y-treated cells appear on the bottom. Scatter plots of **(E)** mitochondrial area and **(F)** mitochondrial lengths in DMSO, and nocodazole-, and noc+Y-treated cells normalised to the mean of the control cells. In B and C, data were obtained from N=3 independent repeats, with >80 cells in each repeat. In **E** and **F**, data were obtained from N=2 independent repeats, with >15 cells in each repeat. The black circles represent the means of the individual repeats, and the black lines represent the mean of the repeats. Asterisks (**) represent p<0.003, and ‘n.s.’ implies no significant difference, 1-way ANOVA with Tukey-Kramer post hoc test.

In the absence of MTs in nocodazole-treated cells, the Rho guanine nucleotide exchange factor GEF-H1 promotes GTP-binding and activation of RhoA, resulting in increased actomyosin contractility^24^. Given a known role for actomyosin contractility in providing pre-constriction to mitochondria to enable Drp1-dependent fission^25,26^, we asked if GEF-H1-dependent actomyosin contractility was responsible for the enhanced mitochondrial fission upon loss of MTs by using the Rho-associated protein kinase inhibitor Y-27632. Cells were treated with DMSO (control), nocodazole, or nocodazole+Y-27632 and their mitochondrial morphologies were visualised (Fig. 2D). Quantification of the mitochondrial morphologies revealed that while nocodazole-treated cells exhibited fragmented mitochondria as before, mitochondria in nocodazole+Y-27632 treated cells did not appear different from those in nocodazole-treated cells, indicating that actomyosin contractility was dispensable for the mitochondrial fragmentation observed in the absence of MTs in these cells (Fig. 2E, F).

### Depletion of microtubule stabilising MAPs induces mitochondrial fission

We then probed if the mitochondrial hyperfission phenotype observed upon treatment with nocodazole could be replicated using depletion of known MT stabilising MAPs. We chose to visualise mitochondria in cells depleted of two well-characterised MAPs, MAP1S and MAP4. MAP1S is a ubiquitously-expressed MAP that is responsible for the maintenance of MTs across cell-cycle stages^27^. MAP4 is another ubiquitous tau-related MAP that promotes MT stabilisation^28^. If the lack of MT association were responsible for mitochondrial fragmentation observed in nocodazole-treated cells, we would expect to observe similar mitochondrial morphologies in cells depleted of key MT stabilising proteins: the decreased MT stability in these cells would result in reduced MT lattice binding sites for mitochondria. Indeed, in cells depleted of MAP1S (Fig. S2C) or MAP4 (Fig. S2D), immunofluorescence images revealed mitochondria that were more fragmented, with smaller areas and lengths, compared to control cells transfected with scrambled siRNA (Fig. 3A-C).

**Figure 3.**
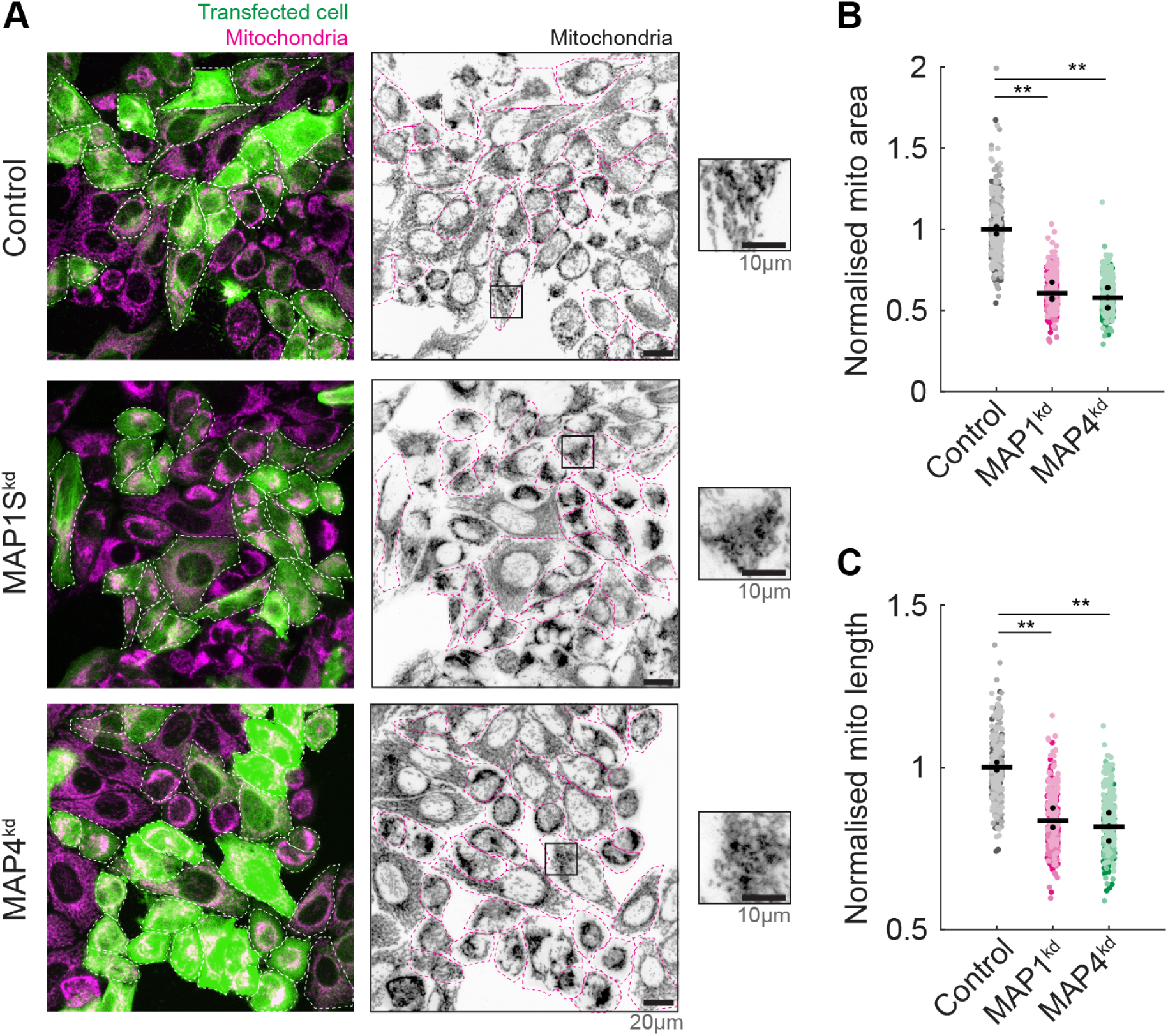
Depletion of MT-stabilising MAPs results in fragmentation of mitochondria. **(A)** Immunofluorescence (IF) images of mitochondria (magenta) in cells transfected (green) with control (top), MAP1S (middle) or MAP4 (bottom) siRNA. The mitochondria channel is represented individually, and the enlarged versions of the regions marked with the black squares from control, MAP1S^kd^, and MAP4^kd^ cells appear on the right. Scatter plots of **(B)** mitochondrial area and **(C)** mitochondrial lengths in control, MAP1S^kd^, and MAP4^kd^ cells normalised to the mean of the control cells. Data were obtained from N=3 independent repeats, with >80 cells in each repeat. The black circles represent the means of the individual repeats, and the black lines represent the mean of the 3 repeats. Asterisks (**) represent p<0.002, 1-way ANOVA with Tukey-Kramer post hoc test.

### Microtubule perturbation alters both mitochondrial form and function

Mitochondrial form is linked to its function, with elongated mitochondrial networks typically linked to more efficient ATP-production, and fragmented mitochondria associated with reduced ATP, but increased ROS production^18^. We tested if these functional differences were apparent in paclitaxel- and nocodazole-treated HeLa cells in two ways. First, we employed the dye TMRE to inspect the mitochondrial membrane potential (functional status) of DMSO (control), paclitaxel-and nocodazole-treated cells. We observed highly reduced mean fluorescence intensity of TMRE in nocodazole-treated cells compared to control cells, whereas paclitaxel-treated cells showed a slight but insignificant elevation in TMRE intensity compared to control cells (Fig. 4A, B).

**Figure 4.**
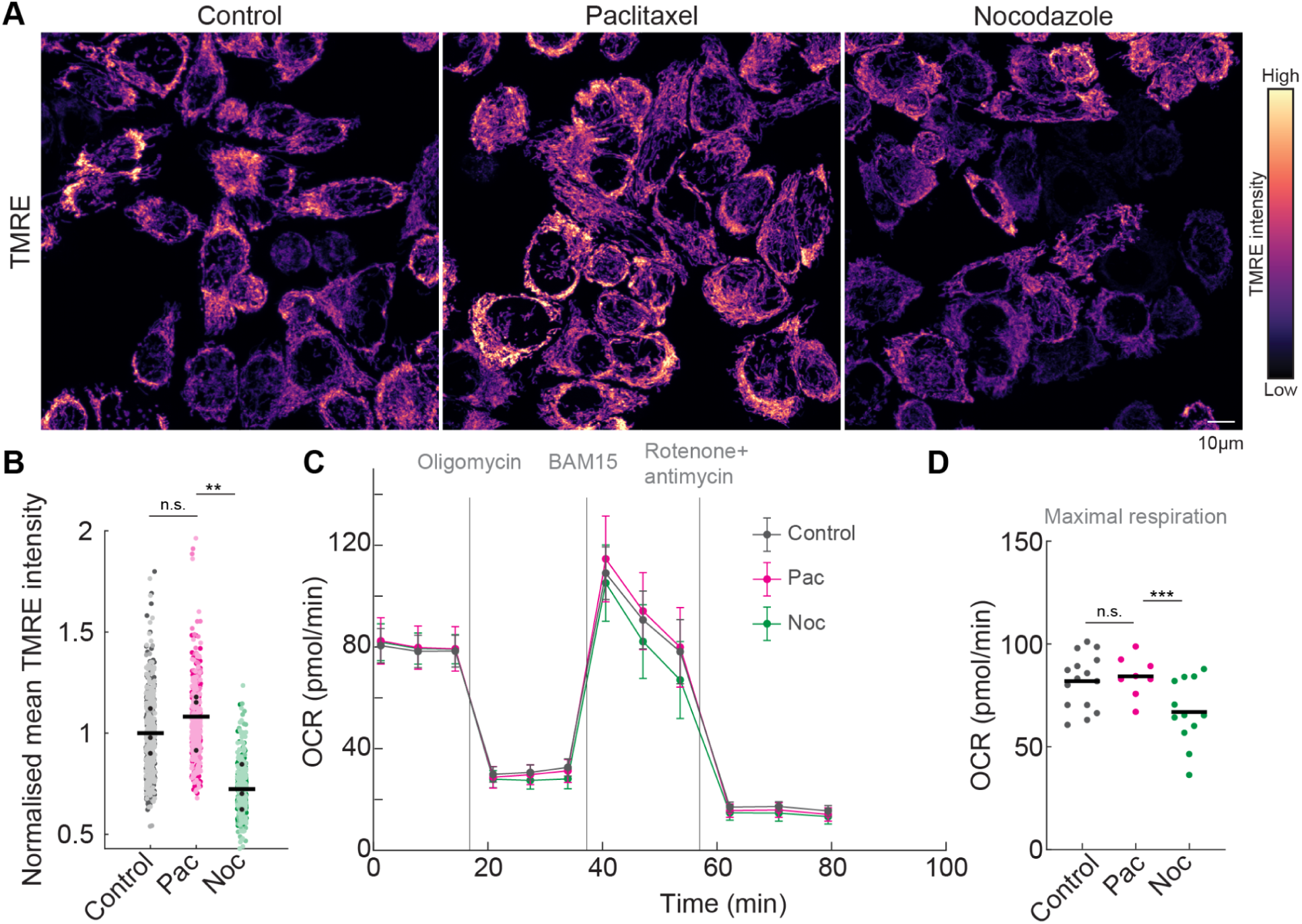
Mitochondrial function is altered upon treatment with nocodazole and paclitaxel. **(A)** Representative live-cell confocal microscopy images of TMRE-stained mitochondria in HeLa cells following treatment with DMSO (control), paclitaxel (pac), and nocodazole (noc) for 1h. **(B)** Scatter plot of the mean TMRE intensity in control, paclitaxel- and nocodazole-treated cells normalised to the mean of the control. Data were obtained from N=3 independent experiments, with >50 cells in each repeat. The black circles represent the means of the individual repeats, and the black lines represent the mean of the 3 repeats. Asterisks (**) represent p<0.03, and ‘n.s.’ implies no significant difference, Kruskal Wallis test for non-parametric data. **(C)** Plot of the oxygen consumption rate (OCR) against time obtained from a Seahorse assay of control, paclitaxel- and nocodazole-treated cells normalised to cell number. **(D)** Maximal respiration upon treatment with BAM15 measured at the 9^th^ timepoint in **C** in control, paclitaxel- and nocodazole-treated cells. Data were obtained from N>8 independent experiments. Asterisks (***) represent p<0.01, and ‘n.s.’ implies no significant difference, 1-way ANOVA with Dunnett’s multiple comparisons test.

Next, we quantified the oxygen consumption rate (OCR) of mitochondria using Seahorse assays. The basal OCR was unchanged in paclitaxel- and nocodazole-treated cells compared to control cells (Fig. 4C). However, the maximal respiration was significantly lowered in nocodazole-treated cells compared to control cells, and paclitaxel-treated cells displayed a minor elevation in maximal OCR (Fig. 4D), mirroring the results we obtained upon quantification of TMRE intensities. Together, nocodazole-treated cells had lowered mitochondrial membrane potential and maximal OCR, consistent with fragmented mitochondria.

### Starvation-induced mitochondrial hyperfusion is preceded by alteration in the underlying microtubule network

Next, we focused on the mitochondrial fusion phenotype that we observed in paclitaxel-treated cells. The mitochondrial morphologies in these cells were reminiscent of mitochondrial hyperfusion that is observed upon nutrient deprivation or starvation in several cell types. This starvation-induced mitochondrial hyperfusion (SIMH) is thought to be a protective response, with the mitochondrial hyperfusion aiding to increase ATP production and preventing mitophagy in these nutrient deprived cells^20^. We simulated starvation in HeLa cells by removing their regular culture medium and incubating them in Hank’s balanced salt solution (HBSS) for 5h as described previously^20,29^. As expected, HBSS-treated cells showed mitochondrial hyperfusion consistent with SIMH (Fig. 5A), that was reflected in the quantification of mitochondrial morphologies (Fig. 5B, C). Surprisingly, we also observed more prominent endogenous tubulin staining in HBSS-treated cells compared to WT control cells (Fig. 5A, D).

**Figure 5.**
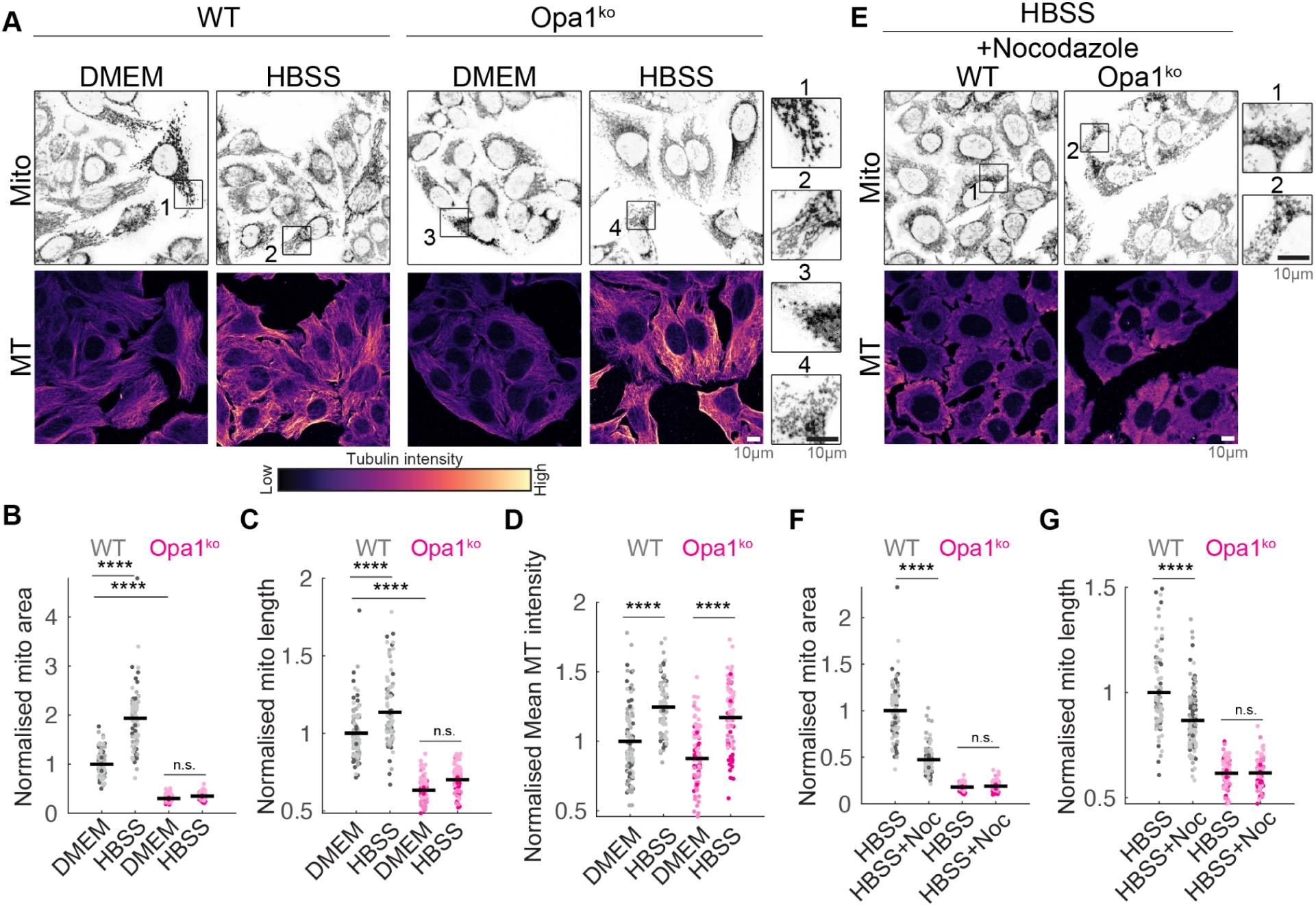
Microtubule network changes maintain mitochondrial hyperfusion in starved cells. **(A)** Representative confocal microscopy images of Mitotracker Deep Red-stained mitochondria (top row), and anti-ɑ-tubulin-stained MT (bottom row) in fixed control (DMEM) and starved (HBSS) WT cells (left), and Opa1^ko^ cells (right). The brightness and contrast of the MT images are set to the same levels to enable comparison. The enlarged versions of the regions marked 1, 2, 3 and 4 from WT-DMEM, WT-HBSS, Opa1^ko^-DMEM, and Opa1^ko^-HBSS cells appear on the right. Scatter plots of **(B)** mitochondrial area, **(C)** mitochondrial lengths and, **(D)** mean MT intensity in WT-DMEM, WT-HBSS, Opa1^ko^-DMEM, and Opa1^ko^-HBSS cells normalised to the mean of the WT-DMEM cells. **(E)** Representative confocal microscopy images of Mitotracker Deep Red-stained mitochondria (top row), and anti-ɑ-tubulin-stained MT (bottom row) in starved (HBSS) WT cells (left), and Opa1^ko^ cells (right) treated with nocodazole. The enlarged versions of the regions marked 1, and 2 from WT and Opa1^ko^ starved cells treated with nocodazole appear on the right. Scatter plots of **(F)** mitochondrial area, and **(G)** mitochondrial lengths in WT-HBSS and Opa1^ko^-HBSS cells treated with nocodazole normalised to the mean of the WT-HBSS cells. In **B**, **C**, **D**, **F** and **G**, data were obtained from N=3 independent repeats, with 20-45 cells in each repeat. The black circles represent the means of the individual repeats, and the black lines represent the mean of the 3 repeats. Asterisks (****) represent p<10^-4^, and ‘n.s.’ implies no significant difference, 2-way ANOVA with Tukey-Kramer post hoc test.

We previously reported in fission yeast that increasing mitochondrial lengths reduces MT depolymerisation rates, and thus enhances MT stability^30^. In the SIMH experiments, we observed simultaneous increase in both mitochondrial length as well as more prominent MT networks. To delineate the effects of mitochondrial size in altering MT prominence, we knocked out the mitochondrial inner membrane fusion protein Opa1 (Opa1^ko^) to create cells with fragmented mitochondria (Fig. S2B, Fig. 5A). In Opa1^ko^ cells, mitochondria appeared fragmented in both control and starved cells due to lack of the fusion protein (Fig. 5A-C). However, the endogenous tubulin intensity in starved Opa1^ko^ cells remained similar to starved WT cells, indicating that starvation induces MT prominence independent, and upstream of mitochondrial hyperfusion.

We next probed if mitochondrial hyperfusion in starved cells was dependent on the underlying MT network by treating nutrient-deprived cells with nocodazole. Indeed, absence of MTs resulted in fragmentation of mitochondria even in starved WT cells (Fig. 5E-G). Taken together, these results indicated that starvation induced changes in the MT network, which were necessary to maintain the enhanced mitochondrial fusion observed in these cells.

### Increased MT association, but not PTMs, underlies mitochondrial hyperfusion in starved cells

The enhanced MT prominence we observed in starved cells could have arisen from increased expression of ɑ-tubulin or increased stability of the underlying MT network. Western blots of ɑ-tubulin in control and HBSS-treated WT and Opa1^ko^ cells revealed no difference in expression, and thus ruled out the first scenario (Fig. 6A).

**Figure 6.**
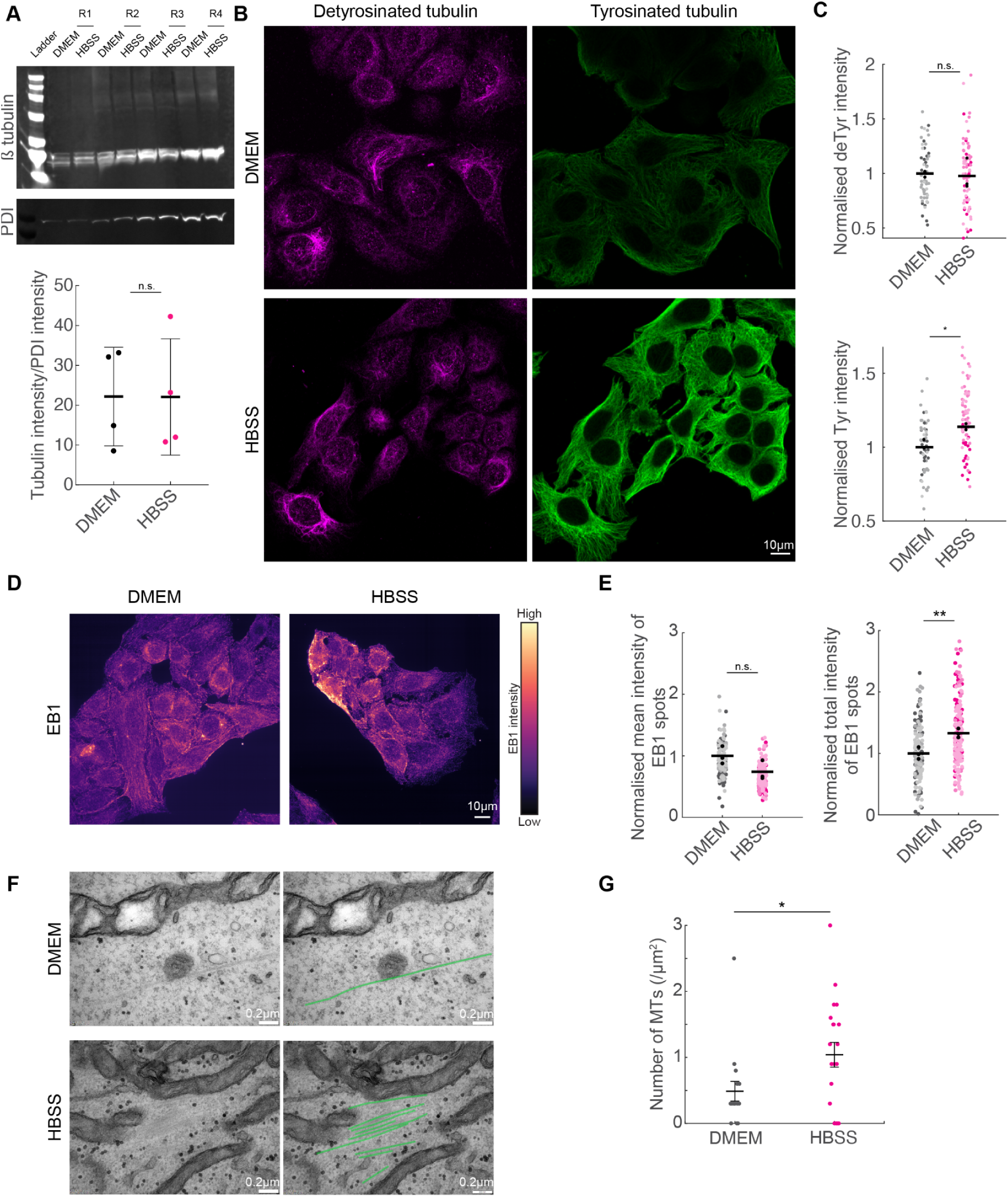
MT numbers, but not detyrosination, are altered during starvation. **(A)** Western blots of β-tublin and PDI in control (DMEM) and starved (HBSS) cells over 4 repeats (R1-4, top), and their quantification (below). **(B)** IF images of detyrosinated tubulin (deTyr, magenta) and tyrosinated tubulin (Tyr, green) in control (DMEM) and starved (HBSS) cells. The brightness and contrast of the images are set to the same levels to enable comparison. **(C)** Scatter plots of deTyr intensities (top), and Tyr intensities (bottom) in DMEM and HBSS cells normalised to the mean of the DMEM cells. **(D)** IF images of EB1 in control (DMEM) and starved (HBSS) cells. **(E)** Scatter plots of mean intensity of EB1 spots (left), and total intensity of EB1 spots (right) in DMEM and HBSS cells normalised to the mean of the DMEM cells. **(F)** TEM images of thin sections of DMEM and HBSS cells (left) with MTs marked in green (right). **(G)** Quantification of the number of MTs per µm^2^ in DMEM and HBSS cells. The asterisk (*) represents p=0.033 (N>15 regions), student’s T-test. In **C** and **E**, data were obtained from N=3 independent repeats, with 15-85 cells in each repeat. The black circles represent the means of the individual repeats, and the black lines represent the mean of the 3 repeats. The asterisk (*) represents p<0.05, asterisks (**) represent p=0.0092 and ‘n.s.’ implies no significant difference, 1-way ANOVA with Tukey-Kramer post hoc test.

Next, we asked if MT stability was enhanced during SIMH. Tubulin acetylation has been used as a marker for stable MT populations, but more recent investigations have revealed the acetylated tubulin instead marks MTs that can withstand compressive/bending forces in cells^31^. Nonetheless, treating control and starved cells with the deacetylation inhibitor trichostatin A (TSA)^32^ did not result in increased mitochondrial fusion, even though tubulin acetylation was significantly enhanced (Fig. S3A-D). Increased detyrosination is a marker of more stable MT populations^33^, and has previously been reported to occur upon starvation in BS-C-1 cells^34^. However, we did not observe increased tubulin detyrosination in starved cells compared to control (Fig. 6B, C).

Strikingly, we observed increased number and intensity of tyrosinated MT populations in starved cells (Fig. 6B, C). This led us to ask if the enhanced prominence of MTs in starved cells resulted from an increase in total MT numbers. Indeed, quantification of the growing MT plus ends using endogenous EB1 staining in control and HBSS-treated cells revealed a significant increase in EB1 intensity in starved cells (Fig. 6D, E). High resolution TEM images confirmed these findings, with starved cells containing ∼2x as many MTs per unit area as cells in full medium (Fig. 6F, G). Thus, starvation appears to stimulate MT polymerisation. This increase in MT mass likely serves to scaffold mitochondria which can then undergo hyperfusion on MT networks.

### Forced association of mitochondria with microtubules results in mitochondrial hyperfusion

From our data in starved cells, we concluded that increasing the available binding sites on MTs for mitochondria led to mitochondrial hyperfusion. To directly test our hypothesis that linking mitochondria to MTs enhances their fusion/reduces their fission, we employed rapamycin-induced heterodimerisation of mitochondrial outer membrane targeted FKBP (mito-FKBP) with MT-bound FRB (MT-FRB) to force the association between MTs and mitochondria. In the absence of rapamycin, and when only mito-FKBP was expressed in cells (with or without addition of rapamycin), we observed normal mitochondrial morphology (Figs. 7A, S4). Upon addition of rapamycin in cells expressing both mito-FKBP and MT-FRB, the mitochondrial networks rapidly underwent fusion such that they eventually resembled starved cells (Fig. 7B-D). To confirm that the forced association of mitochondria to MTs in these experiments led to increased fusion, rather than close apposition, we visualised mitochondria in TEM images. Indeed, in the presence of rapamycin in mito-FKBP+MT-FRB cells, mitochondria appeared fused and fully contiguous (Fig. 7E). Taken together, we demonstrate that association of mitochondria with microtubules is sufficient to result in hyperfused networks.

**Figure 7.**
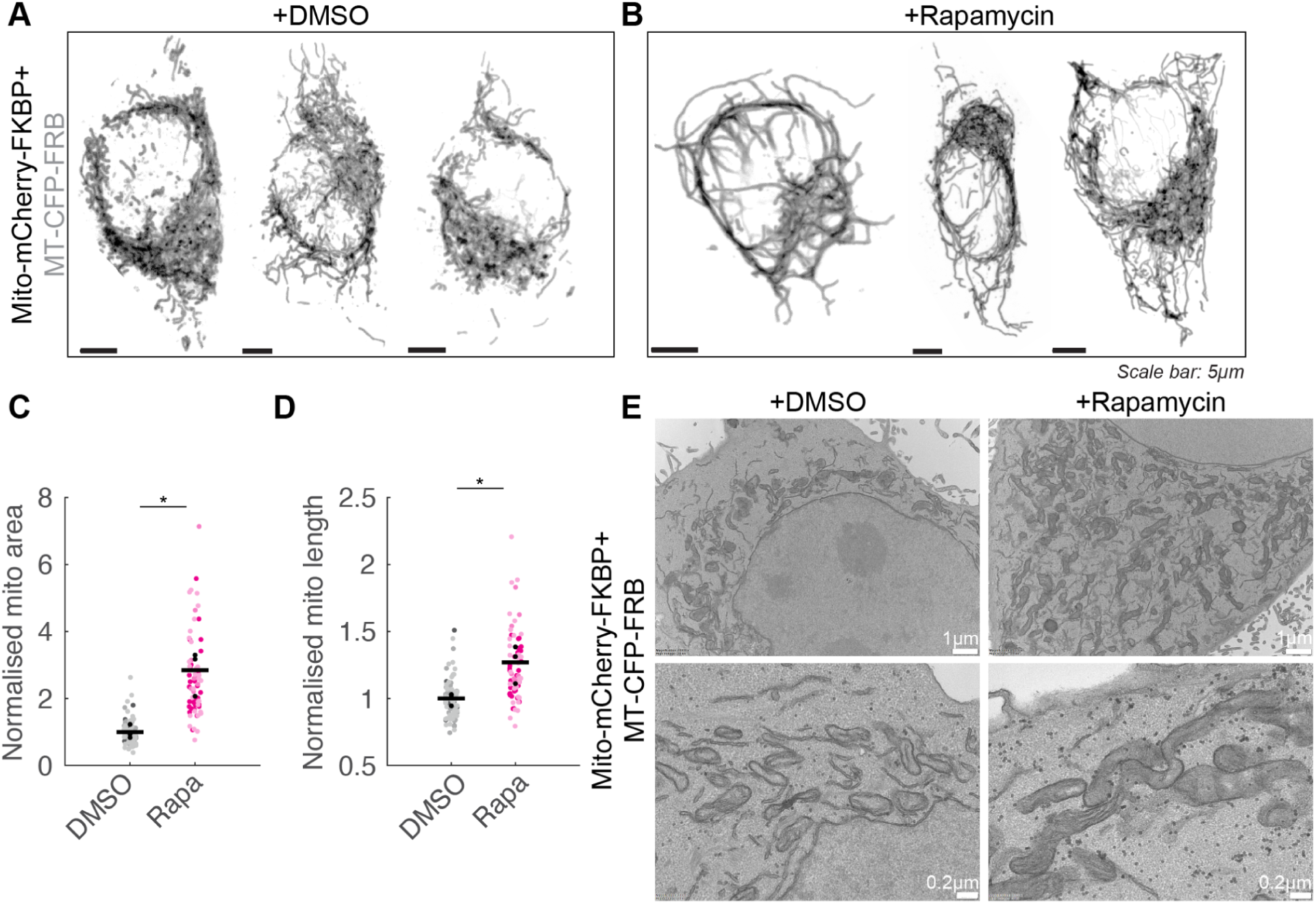
Forced association of mitochondria to microtubules leads to mitochondrial hyperfusion. **(A)** Representative confocal microscopy images of mitochondria in cells transfected with mito-mCherry-FKBP and MT-CFP-FRB in control (DMSO) and **(B)** rapamycin-treated cells. Scatter plots of **(C)** mitochondrial area and **(D)** mitochondrial lengths in control (DMSO) and rapamycin-treated cells normalised to the mean of the control cells. Data were obtained from N=3 independent repeats, with 10-45 cells in each repeat. The black circles represent the means of the individual repeats, and the black lines represent the mean of the 3 repeats. The asterisk (*) represents p<0.05, 1-way ANOVA with Tukey-Kramer post hoc test. **(E)** TEM of thin sections of control (left) and rapamycin-treated cells (right) at low (top) and high (bottom) magnifications, showing contiguous, long mitochondria in the rapamycin-treated cells.

## Discussion

Mitochondrial fission and fusion have a central role in determining cellular function and thus have been the subject of investigation for decades. Mitochondria have long been known to associate with MTs to regulate their distribution^2^. Here, we discovered that association with MTs additionally controls mitochondrial fission-fusion status (Fig. 1). Similar to our findings in fission yeast^8,10^, depolymerisation of MTs led to mitochondrial morphologies consistent with increased fragmentation, while stabilisation of MTs resulted in longer mitochondria. Given the absence of mitochondrial fragmentation in Drp1^ko^ cells treated with nocodazole (Fig. 2A-C), we concluded that the lack of MT association enhances Drp1-driven mitochondrial fission. Indeed, in fission yeast, mitochondrial fission is inhibited when mitochondria are bound to MTs due to the inability of Dnm1 (yeast Drp1) to assemble around MT-bound mitochondria^8^. However, loss of underlying microtubule tethers could also reduce mitochondrial fusion rates by decreasing both the frequency and stability of mitochondrial encounters required for fusion.

Our findings are consistent with and complementary to several previous studies examining the relationship between cytoskeletal dynamics and mitochondrial morphology. While a previous report demonstrated that reduction of mitochondrial membrane tension by nocodazole treatment reduces mitochondrial fission rates^35^, mitochondrial morphology was not quantified prior to fission rate measurements in that study; from the images presented, mitochondria in nocodazole-treated cells appear smaller than in controls, suggesting that earlier time point measurements may have yielded different outcomes. Similarly, actin ‘waves’, cycles of actin polymerisation and depolymerisation that promote mitochondrial mixing in interphase cells, have been shown to drive fission of actin-enveloped mitochondria^36^, with nocodazole appearing to promote interconnectedness of actin-enveloped mitochondria rather than the fragmentation we observe. The increased mitochondrial fragmentation in our experiments likely reflects the net outcome of altered fission dynamics and reduced fusion over our specific treatment conditions, where impaired MT-dependent mitochondrial positioning and reduced fusion may dominate over any wave-mediated fission pathway. This interpretation is supported by the observation that inhibition of actomyosin contractility with the ROCK inhibitor Y-27632 did not reverse the fragmentation phenotype (Fig. 2D-F), suggesting that ER tubule-mediated pre-constriction via actomyosin^37^ is not the primary driver of the Drp1-dependent fission observed under these conditions.

In this work, we also tested the general effect of loss of microtubule stability and reduced mitochondrial association in regulating mitochondrial dynamics in cells depleted of MAP1S and MAP4, and observed mitochondrial fragmentation phenotypes (Fig. 3). This data is similar to the mitochondrial morphology seen in nocodazole-treated cells and is expected given the loss of MT stability and thus reduced mitochondria-MT association in the MAP-depleted cells. Differential MAP decoration is known to alter motor loading, and thereby, cargo transport and positioning on MTs^38^. So too, depletion of motor proteins has been observed to alter mitochondrial dynamics on MTs^39,40^. Thus, it would be important in future to test motor-dependent effects of mitochondrial positioning on MTs, and how this might result in altered mitochondrial dynamics.

Mitochondrial form is linked to its function, with fragmented mitochondria typically producing low ATP and high ROS^18^. As reported previously in other cell types, we observed reduced mitochondrial membrane potential and maximal respiration in fragmented, nocodazole-treated cells^41^. Paclitaxel-treated cells also showed a slight but insignificant increase in mitochondrial membrane potential and maximal OCR (Fig. 4).

SIMH acts as a survival mechanism to counter mitophagy and to enable efficient ATP production via the fused mitochondria^20^. Here, we revealed that an increase in MT prominence occurs independently of the hyperfusion phenotype (Fig. 5A-C). We previously showed that increasing mitochondrial association with MTs by altering mitochondrial form could reduce MT depolymerisation rates^30^. To establish the temporal sequence of mitochondrial hyperfusion and increased MT prominence observed in starved cells, we employed Opa1^ko^ cells that have small, fragmented mitochondria due to unopposed mitochondrial fission. Mitochondrial hyperfusion following nutrient deprivation has been shown previously to result from down-regulation of Drp1^20^, or dependent on mitochondrial fusion proteins Mfn1 and Opa1^42^. Here, we observed that while mitochondria continued to remain fragmented even in starved Opa1^ko^ cells, the MT prominence was unaltered in these cells.

The increased MT prominence we observed in our starved and fixed cells stained for tubulin could have arisen from increase in tubulin expression, enhanced MT stability, or higher overall tubulin expression. We ruled out an increase in tubulin expression since comparison of Western blots from control and starved cells did not show any difference (Fig. 6A). Tubulin post translational modifications (PTMs) enable diversification of MT functions such as stability, primarily after they have been polymerised into MTs^33,43^. Previous studies have reported increased tubulin PTMs acetylation^44^ or detyrosination^34^ in nutrient-deprived cells. Tubulin acetylation was originally thought to be a marker of stable MT populations, however, more recent data suggests that acetylation enhances MTs’ ability to withstand compression in cells^31^. Tubulin detyrosination is the key PTM that is associated with stable MTs, while tubulin tyrosination marks labile populations^45^. We did not observe an increase in tubulin acetylation upon starvation in our system. While we did not observe changes in the detyrosinated microtubule population, we saw a marked increase in the population of tyrosinated MTs (Fig. 6B, C). Thus, we tested if the MT prominence in starved cells resulted from an increase in MT numbers by quantifying EB1 levels (Fig. 6D) and counting MT numbers in EM thin sections (Fig. 6F). Indeed, we observed a significant increase in EB1 levels and total MT numbers in starved cells (Fig. 6E, G). Our high resolution EM images also suggest that MTs might undergo more bundling under starvation (Fig. 6F). This enhancement in MT numbers likely provides an increase in MT lattice regions for mitochondrial association thus promoting docking and increased fusion of MT on microtubules.

The upstream signal connecting nutrient deprivation to increased MT polymerisation remains an open question. The canonical nutrient sensors AMPK and mTORC1 are activated and inhibited by starvation respectively, yet neither has been shown to have a straightforward relationship with MT polymerisation thus far. While AMPK phosphorylates the MT plus-end protein CLIP-170 to promote MT dynamics^46^, its effects on MT accumulation are context-dependent. Interestingly, yeast TORC1 has been shown to interact with EB1 and CLIP-170 via its Raptor subunit, and TORC1 inhibition results in hyperelongated MTs^47^. Reduction in mTORC1 activity upon starvation could potentially promote MT polymerisation and contribute to the increased EB1 levels we observed. While mTORC2’s cytoskeletal role is primarily rapamycin-insensitive and actin-focused^48^, identifying the signal that connects nutrient status to MT polymerization, and how this feeds into mitochondrial hyperfusion as a survival response represents an important avenue for future investigation.

MTs play a key role in controlling mitochondrial position and transport in mammalian cells. Motor proteins dynein and kinesin and their adaptors TRAK1 and TRAK2 mediate the minus and plus end-directed movement respectively of mitochondria on MT tracks^2^. In neurons, TRAK1 glycosylation at regions of high glucose^49^ as well as syntaphilin-kinesin coupling in a Ca^2+^-dependent manner enable docking of mitochondria on MTs^13,14^. Here, we demonstrated that forcing the association between mitochondria and MTs in non-neuronal cells is sufficient to induce extensive mitochondrial fusion (Fig. 7). Future studies will be invaluable to reveal if there is a novel anchor for mitochondria on MTs, that could be leveraged to manipulate mitochondrial form and function.

Taken together, these findings raise important questions for future investigation: What are the molecular anchors that tether mitochondria to MTs across cell types? How does MT association mechanistically regulate fusion through encounter frequencies, membrane mechanics, or other pathways? Can manipulation of mitochondrial-MT interactions be leveraged therapeutically in diseases characterised by dysregulated mitochondrial dynamics? Addressing these questions will be essential for understanding how cells coordinate mitochondrial form with function and may reveal new strategies for manipulating mitochondrial networks in metabolic and neurodegenerative diseases.

## Acknowledgements

We thank M. Ahmed for constructive comments on the manuscript and D. Shu (Optometry, UNSW) for technical assistance. We acknowledge the facilities and the scientific and technical assistance of the Katharina Gaus Light Microscopy Facility (KGLMF) at UNSW Sydney. VA is supported by EMBLA Australia Funding; VA and GMIR acknowledge funding from the Australian Research Council Discovery Projects Grant (DP260100749).

