## Supplementary Information for "Mitochondrial morphology and function are tuned by microtubule association"

### Supplementary Figures

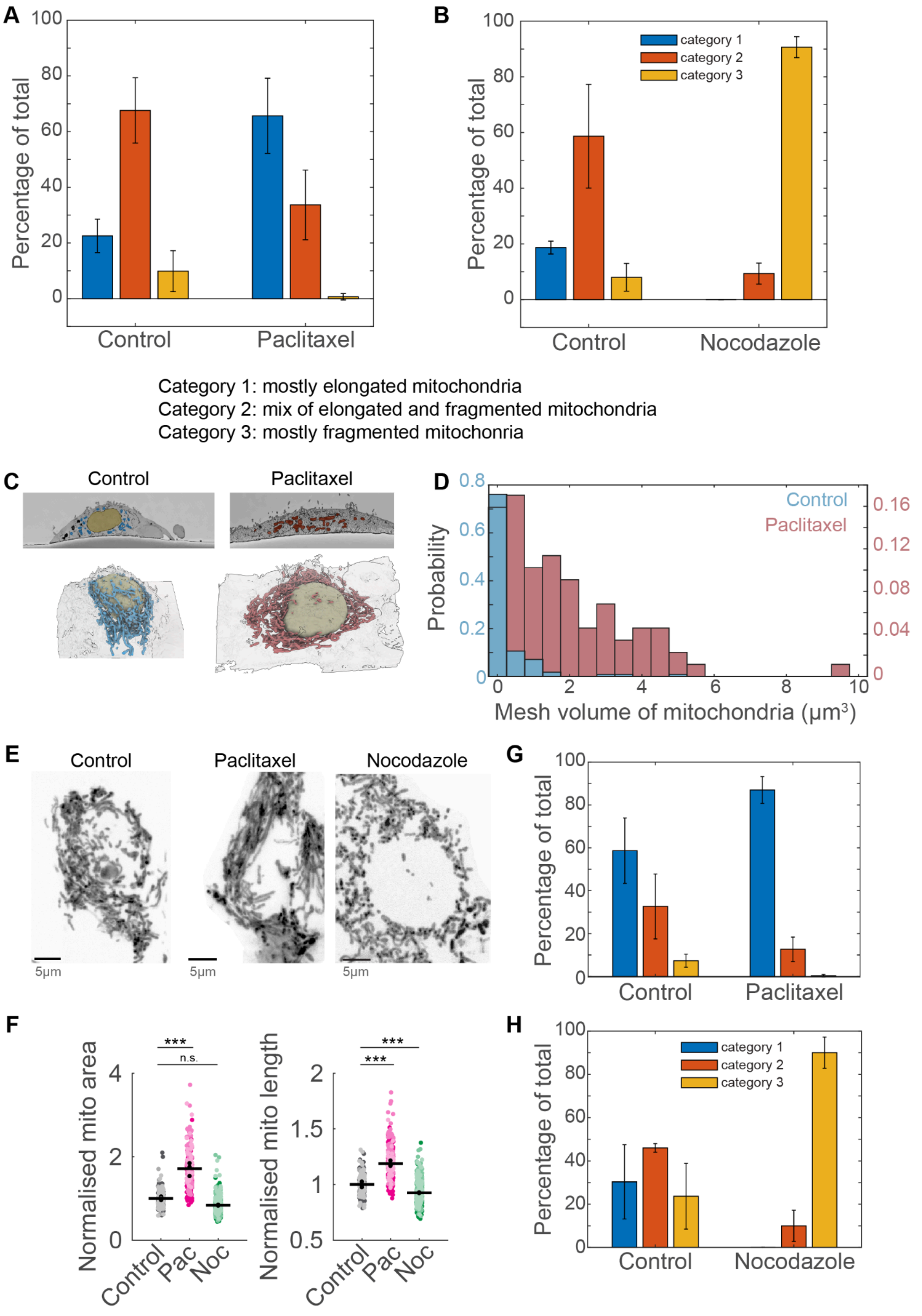


**Figure S1. Microtubule perturbation alters mitochondrial form in HEK293 cells.**
Classification of HeLa cells into three categories listed below the figure based on mitochondrial morphology in **(A)** DMSO and paclitaxel-treated cells and **(B)** DMSO and nocodazole-treated cells. **(C)** SBF-SEM images of control and paclitaxel-treated cells. **(D)** Histogram of mitochondrial volume in control and paclitaxel-treated cells showing a high proportion of

mitochondria with substantially increased volumes in the paclitaxel-treated cell compared to the control cell. **(E)** Representative live-cell confocal microscopy images of Mitotracker Deep Red-labelled mitochondria in HEK293 cells following treatment with DMSO (control), paclitaxel, and nocodazole for 1h. **(F)** Scatter plots of mitochondrial area (left) and mitochondrial lengths (right) in control, paclitaxel- and nocodazole-treated cells normalised to the mean of the control cells. Data were obtained from N=3 independent repeats, with 25-115 cells in each repeat. The black circles represent the means of the individual repeats, and the black lines represent the mean of the 3 repeats. Asterisks (***) represent $p<0.001$, and 'n.s.' implies no significant difference, 1-way ANOVA with Tukey-Kramer post hoc test. Classification of HEK293 cells into three categories listed in panel **A** based on mitochondrial morphology in **(G)** DMSO and paclitaxel-treated cells and **(H)** DMSO and nocodazole-treated cells.

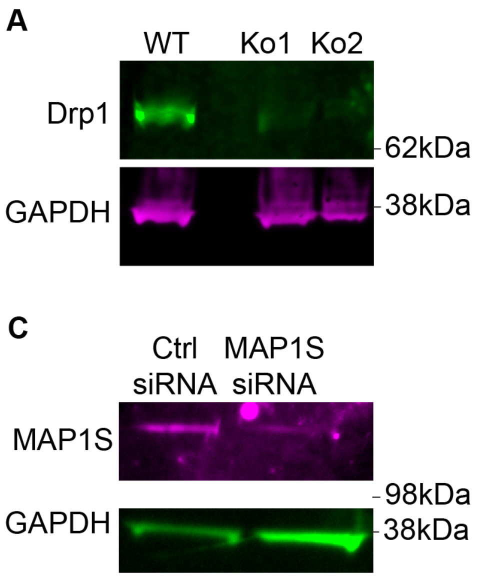


**Figure S2. Confirmation of Drp1 and Opa1 knockouts, MAP1S and MAP4 depletion using Western blots**

Western blots confirming knockout of **(A)** Drp1, **(B)** Opa1 and depletion of **(C)** MAP1S and **(D)** MAP4 in HeLa cells. GAPDH was used as loading control in all cases.

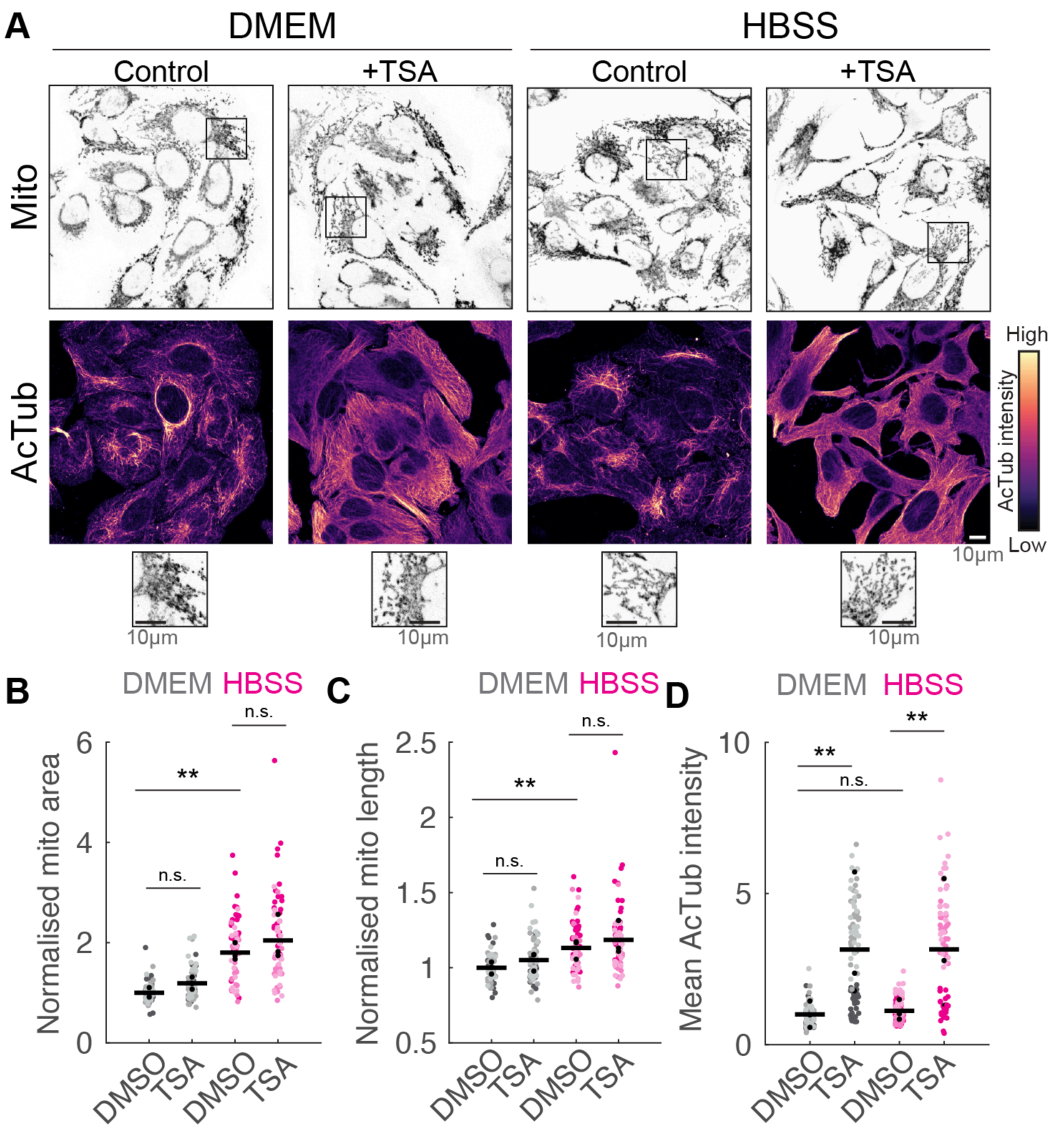


**Figure S3. Tubulin acetylation is not enhanced in starved cells.**

**(A)** IF images of acetylated tubulin (AcTub) in Mitotracker Deep Red-stained control (DMEM) and starved (HBSS) cells in the absence and presence of TSA. The brightness and contrast of the AcTub images are set to the same levels to enable comparison. The enlarged versions of the regions marked with the black square appear below the images. Scatter plots of **(B)** mitochondrial area and **(C)** mitochondrial lengths in control (DMSO), and TSA-treated control (DMEM) and starved (HBSS) cells normalised to the mean of the control cells (DMSO-treated WT cells). **(D)** Scatter plot of mean AcTub intensities in control (DMSO), and TSA-treated DMEM and HBSS cells normalised to the mean of the control cells (DMSO-treated DMEM cells). In **B**, **C** and **D**, data were obtained from N=3 independent repeats, with >22 cells in each repeat. The black circles represent the means of the individual repeats, and the black lines represent the mean of the 3 repeats. Asterisks (**) represent $p<0.01$, and 'n.s.' implies no significant difference, 2-way ANOVA with Tukey-Kramer post hoc test.

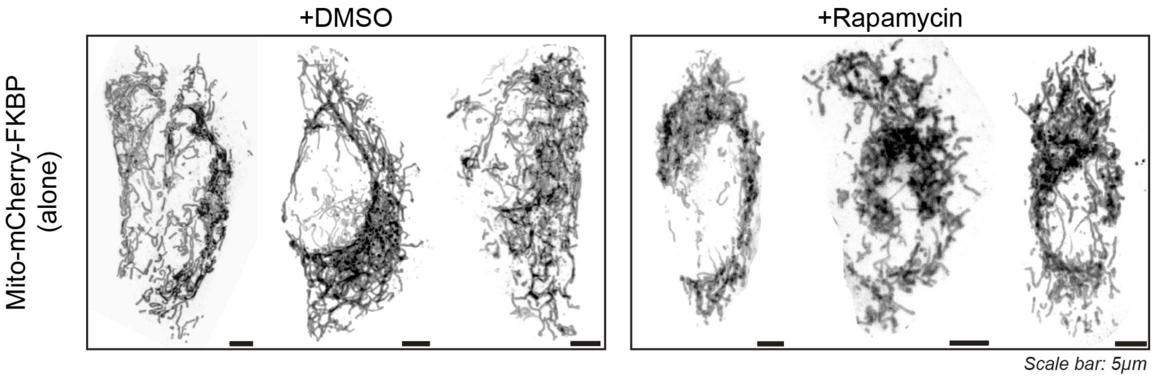


**Figure S4. Mitochondrial network in cells expressing only mito-FKBP**
Representative confocal microscopy images of mitochondria in cells transfected with mito-mCherry-FKBP alone in control (DMSO, left) and rapamycin-treated (right) cells.
